# Latitudinal effects on weight and age-dependent survival of red fox *Vulpes vulpes*

**DOI:** 10.64898/2026.06.29.735157

**Authors:** Tomas Willebrand, Morten Odden, Gustaf Samelius, Zea Walton, Kjartat Østbye, Jan Englund, Göran Spong

## Abstract

Age-dependent survival is central to understanding population dynamics and life-history evolution. We analysed carcass weight and age-at-harvest data from 6 022 red foxes (*Vulpes vulpes*) collected across Sweden between 1967 and 1971 to evaluate latitudinal effects on body mass and age-dependent survival. Carcass weights decreased from south to north in both adults and sub-adults, contrary to Bergmann’s rule, with southern foxes weighing approximately 1.27 times more than northern foxes. The latitudinal weight gradient exceeded the sex difference in both age classes, and no sex *×* region interaction was detected. The decrease in weight with latitude is consistent with reduced prey availability and harsher winter conditions in the north, which limit growth and body size during development. Using a Bayesian age-at-harvest model with region-specific population growth rates (*λ*), we estimated age-dependent survival probabilities for four latitudinal regions and both sexes. Despite the strong latitudinal gradient in weight, survival did not show a corresponding pattern — regional differences were uncertain, with all credible intervals spanning zero. Regional population growth rates were consistent with slight decline in the north and near-stability in the south-central region, which suggests that body condition and population dynamics are coupled at the regional scale despite no survival gradient. The decoupling of body condition and survival across regions suggests that mortality patterns are similar across the latitudinal gradient. We discuss these patterns in terms of latitudinal productivity gradients, prey availability, and life-history trade-offs in a widely distributed carnivore.

## Introduction

Body size is a key trait that varies both within and between species (Blanckenhorn, 2000), and plays a central role in the theory of life-history evolution (Stearns, 2000; Clauset and Erwin, 2008). It is a major determinant of metabolic rate, resource requirements, and ecological interactions (Brown et al. 2004), and is closely linked to other life-history traits such as reproduction and survival (Roff, 1986; Sibly and Brown, 2007; Brooks et al. 2016). Body size also shows consistent geographic variation in many species, often following latitudinal gradients (Blanckenhorn, 2000; Huston and Wolverton, 2011), which has been the subject of much research and debate in ecology and evolution (McNab, 1971; Blackburn et al. 1999). Understanding the drivers of body size variation and its consequences for population dynamics is therefore important. In general, body size is determined by resource availability during growth (Geist, 1987), and reduced food availability affects body size in high density deer populations (Albon et al. 1987; Festa-Bianchet et al. 2000; Gaillard, Loison, et al. 2003).

Species with a wide geographical distribution are exposed to a broad range of resource availability and climate conditions, which potentially could result in different optimal values of life history-traits. However, as long as the different subpopulations are connected, evolution should favor adaptive phenotypic plasticity (McNamara and Houston, 1996; Kawecki and Stearns, 1993). For example, Meiri, Dayan, et al. (2009) found limited evidence of a body size gradient from the core part to the edge of carnivore distribution, and proposed that the lack of a gradient was due to high dispersal rates that maintain gene flow across the distribution. Furthermore, shifts in age structure and age-dependent survival may also affect body size, as individuals that survive to older ages may grow larger than those that die at younger ages (Theriot et al. 2024).

Patterns of age dependent survival are species specific (Martin, 1995; Loison et al. 1999), and determine resource allocation strategies as body size and morphology (Roff, 1986; Sibly and Brown, 2007; Brooks et al. 2016). Life-history traits such as longevity, age at maturity, and reproductive rate show positive covariation that results in a slow-fast continuum of population dynamics (Oli, 2004), and is closely correlated to generation time (Kraus et al. 2005). Slow species are long-lived with low reproductive rates, whereas fast species are short-lived and allocate resources to early maturation and high fertility; in addition, the effects of stochastic processes on population dynamics are higher at the fast end of the continuum (Sæther et al. 2013).

The red fox *Vulpes vulpes* is a highly adaptive species, and one of the most widely distributed carnivores (Larivière and Pasitschniak-Arts, 1996). It can utilize a wide range prey from voles to roe deer (Englund, 1970; Kjellander and Nordström, 2003), and also feed on berries and fruit as well as garbage (Carvalho and Gomes, 2001). It benefits from human settlements and agricultural activities (Jahren et al. 2020; Gallant et al. 2020; Vuorisalo et al. 2014), but is limited by deep snow and harsh winters at northern latitudes (Bartoń and Zalewski, 2007; Frafjord and Stevy, 1998). Demographic data and other life-history characteristics place the red fox on the fast side of the fast-slow continuum (Heppell et al. 2000). Females start to reproduce at an age about 10 months (Englund, 1970), and survival estimates are low both for adults and subadults (Devenish-Nelson et al. 2013; Willebrand et al. 2022). Despite extensive mortality during the first outbreaks of sarcoptic mange, red fox populations appeared to recover within decades after the outbreak (Soulsbury et al. 2007; Willebrand et al. 2022).

The red fox is distributed throughout all of Sweden’s 450,295 km^2^, which stretches from 55.38^◦^ to 67.86^◦^ latitude. The northern parts are sparsely populated, and the area north of the polar circle (66.34^◦^). The southern parts, on the other hand, have an animal diversity and human densities similar to central European countries. The aim with this study is to evaluate the effects of a latitudinal gradient on age related body size of males and females, and estimate age-dependent survival patterns of red foxes in Sweden. We expect that both body size and survival will decrease with latitude due to lower prey availability and harsher climate conditions in the north. We also expect that the age structure of the population will differ between regions, with a higher proportion of younger individuals in the north due to lower survival rates.

## Methods

### Collecting harvested foxes

The data used in this analysis was part of a monitoring program that was initiated by the late professor Englund to investigate age, reproduction and morphological characteristics of the red fox. In this study, we used 6 022 harvested foxes that were collected from hunters between 1967 and 1971 when efforts were made to retrieve foxes from all parts of Sweden. Carcass weight was recorded to the nearest hectogram, and age was determined by microscopic examination of cementum layers of the canine teeth. See Englund (1965) and Englund (1970) for additional details.

### Latitudinal regions

We grouped the collected foxes in four latitudinal regions similar to (Englund, 1970). See (Zheng et al. 2004; *Geography of Sweden* 2025; *Markanvändningen i Sverige =* 2013; Englund, 2019; Englund, 2020) for more details. Sweden covers a wide range of climatic zones, and stretches more than 1 500 km from south to north. National harvest statistics (*Viltdata* 2025) on red foxes were used to estimate indices of population change.

**N:** *1461 males and 1128 females.* The northernmost part of Sweden that stretches mostly north of 62^◦^ latitude to north of the polar circle. The region is characterized by low productivity and a net primary production (NPP) below 300. The region is dominated by boreal forests (68%) and open tundra (27%), and experienced long-winters and deep snow cover during the study. Microtines *Arvicolinae*, mountain hares *Lepus timidus* and grouse *Tetraoninae* are common prey species for red foxes. Garbage, slaughter remains and carcasses from domestic reindeer *Rangifer tarandus* and moose *Alces alces* are occasionally available in autumn and winter. The region has less than 6 inhabitants per km^2^.
**NC:** *553 males and 437 females.* North-Central Sweden, between 60^◦^ and 62^◦^latitudes, is dominated by boreal conifer forest (84%) interspersed with agricultural (4%) areas. The northwest parts contain open tundra (7%). Winters vary, and snow cover usually stayed from November until March during the study. Microtines, mountain hares and grouse but also roe deer *Capreolus capreolus* are among important prey species. Garbage, slaughter remains and carcasses from moose are occasionally available in autumn and winter. Human settlements are more common than in N, and the region has about 12 inhabitants per km^2^.
**SC:** *929 males and 809 females.* South-Central Sweden, between 58^◦^ and 60^◦^latitudes is characterized by forests (63%) and agriculture (22%) areas. Broad leaf tree species are common in forests. Winters are short with low amounts of snow. Prey species are more diverse than in NC but microtines are still important. Garbage and slaughter remains from ungulate species are available in autumn and winter. The region has among the highest population densities in Sweden, with close to 100 inhabitants per km^2^, and 6% is classified as urban areas.
**S:** *374 males and 289 females.* Southern part of Sweden, which is below 58^◦^ latitude. The region contains forest (65%) and agriculture (19%) areas, and forests contain several different broad leave species in addition to conifers. It is dominated by short winters and a climate similar to central Europe. Prey is abundant, including rabbits *Oryctolagus cuniculus* that are not present in regions further north (Erlinge et al. 1984). Garbage and slaughter remains are available occasionally in autumn and winter. Inhabitants are close to 100 per km^2^ and 7% is classified as urban areas.

### Analysis

We evaluated the effect of region, sex and age on carcass weight using generalized linear models with a Gamma distribution and log link, which was preferred over a Gaussian model based on AIC and residual diagnostics. The Gamma distribution naturally accommodates the increase in variance with predicted weight that is characteristic of body mass data. We separately analysed adults (age *≥* 1 year) and sub-adults (age 1). For adults, age was included as a continuous covariate to account for continued weight gain beyond the first year. Thus, sub-adults represent a single first-year age class where growth is substantially higher compared to adults, so age was not included in the sub-adult model. Note that within-class age variation was not recorded, making the two models non-comparable in structure. Region and sex were treated as categorical variables. We tested for a sex *×* region interaction in both models by comparing additive and interaction models using AIC. We used residual diagnostics to validate models with the package DHARMa (Hartig, 2022).

Age-at-harvest data are routinely collected for many wildlife species and can be used to supplement survival estimates from mark-recapture and known-fate studies. Life-table analysis based on age at harvest data is an indirect method to estimate survival, and its use is restricted by assumptions of a stable age structure, constant survival, and a population at equilibrium (*λ* = 1). These assumptions have been evaluated and to different degrees relaxed (Udevitz and Ballachey, 1998), and a recent study by Skelly et al. (2023) proposed a Bayesian approach to estimate survival probability from age-at-harvest data when auxiliary data to validate assumptions are not available. The model includes flexible definitions of the likelihood and can account for uncertainty in age structure, population growth rate, and harvest selectivity by assigning prior distributions. In addition, survival probability may vary as a function of environmental variables.

Here we have used the outlined Bayesian model proposed by Skelly et al. (2023). Our data contained age classes from first year sub-adults (age-class 1) to 12 years of age. We pooled all ages above 7 years to age class 7+, and tabulated data by age, sex and region. We did not include any parameter on the relative selectivity to be harvested. See section Appendix. We developed 8 models with different combinations of sex, age and region affecting survival probabilities, listed in Table 3. Model fit was evaluated with posterior predictive checks (Kéry, 2010), and used deviance information criterion (*DIC*) for model selection. All models were developed in a Bayesian framework using the JAGS-language (Plummer et al. 2003) (see appendix). All data preparation and analysis were made in R (R Core Team, 2024).

## Results

### Weight decrease with latitudes

Carcass weights of red foxes decreased consistently from the southernmost to the northernmost region in both adults and sub-adults, and for both sexes (Figure 1, Tables 1 & 2). Gamma models with a log link were strongly preferred over Gaussian models in both age classes (ΔAIC *>* 32 for sub-adults, *>* 113 for adults), and residual diagnostics confirmed that heteroscedasticity was resolved under the Gamma distribution (Levene test non-significant for both models). A sex *×* region interaction was not supported in either age class (ΔAIC = *−*1.1 for adults; 0.16 for sub-adults), indicating that the sex difference in weight was consistent across all four regions.

**Figure 1:**
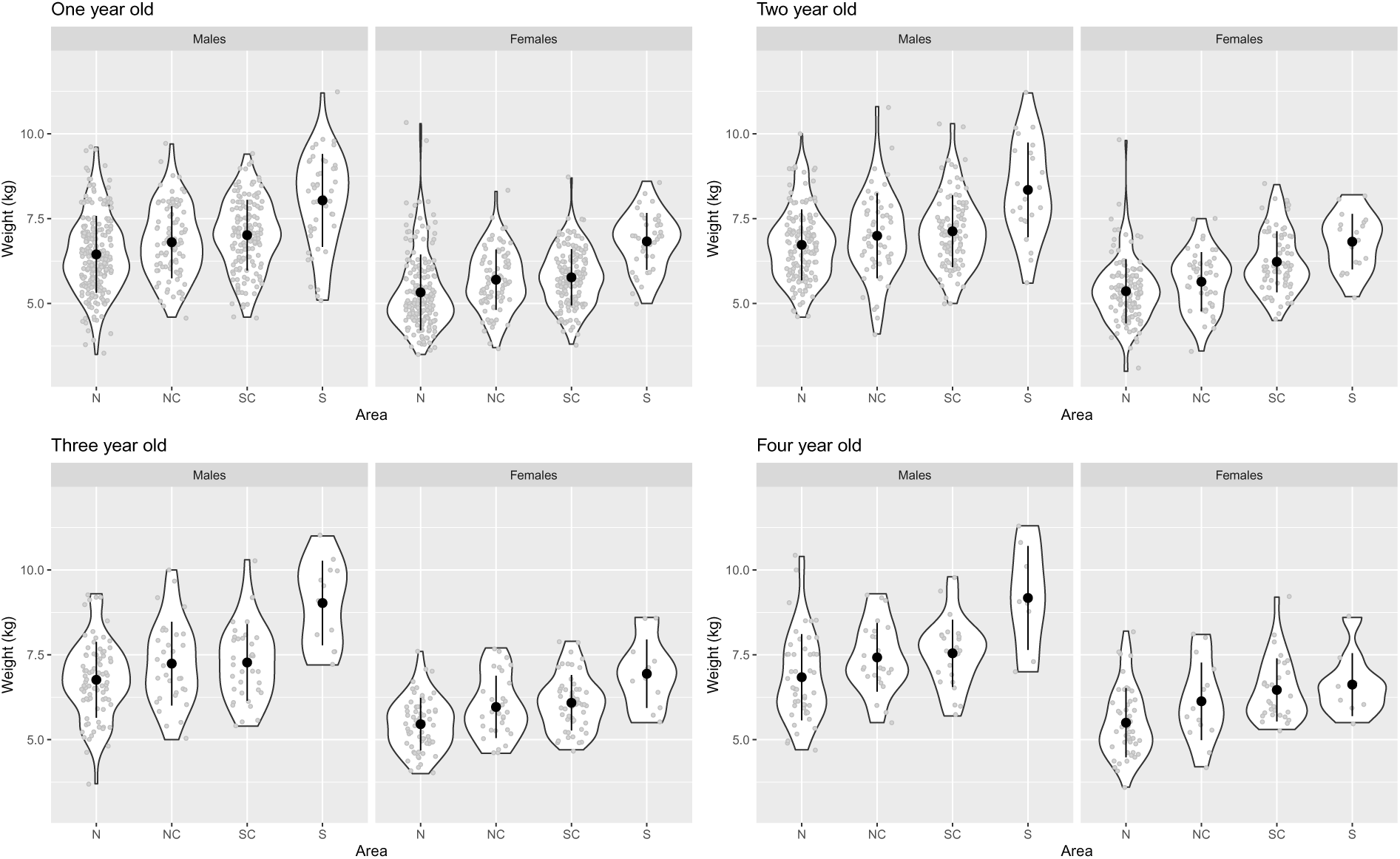
Distribution of carcass weights of foxes aged 1-4 years in a latitudinal gradient of four regions in Sweden.

In adults, foxes in the South were on average 1.27 times heavier than foxes in the North (exp(0.239) = 1.27; Table 1), and males were on average 1.21 times heavier than females (exp(0.190) = 1.21). The intermediate regions followed a clear latitudinal gradient, with North Central and South Central significantly heavier than the North reference. Adults also increased in weight with age, gaining approximately 1.1% per year (exp(0.011) = 1.011), a small but significant effect relative to the regional and sex differences.

**Table 1:** Evaluation of variation in weight dependent on sex, age and region on adult red foxes in Sweden. A generalised linear model with Gamma distribution and log link was used to model the variation. Coefficients are on the log scale; exp(coefficient) gives the multiplicative effect on weight in kg. Numbers in parenthesis are standard errors of the parameter.

|  | *Coefficient (log scale)* | *Multiplier* |
| --- | --- | --- |
| Age | 0.011***<br>(0.002) | 1.011 |
| North Central | 0.067***<br>(0.009) | 1.069 |
| South Central | 0.093***<br>(0.008) | 1.097 |
| South | 0.239***<br>(0.013) | 1.270 |
| Female | -0.190***<br>(0.007) | 0.827 |
| Intercept | 1.858***<br>(0.008) |  |
| Observations | 2,343 |  |
| AIC | 6,794 |  |
| *Note:* | *p<0.1; **p<0.05; ***p<0.01 |  |

In sub-adults, the latitudinal gradient was equally pronounced: foxes in the South were 1.28 times heavier than those in the North (exp(0.243) = 1.28; Table 2), and males were 1.19 times heavier than females (exp(0.174) = 1.19). The latitudinal effect on weight therefore exceeded the sex difference in both age classes, and the pattern was fully consistent across males and females.

**Table 2:** Evaluation of variation in weight dependent on sex and region for sub-adult red foxes in Sweden. A generalised linear model with Gamma distribution and log link was used to model the variation. Coefficients are on the log scale; exp(coefficient) gives the multiplicative effect on weight in kg. Numbers in parenthesis are standard errors of the parameter.

|  | *Coefficient (log scale)* | *Multiplier* |
| --- | --- | --- |
| North Central | 0.037***<br>(0.009) | 1.037 |
| South Central | 0.069***<br>(0.008) | 1.072 |
| South | 0.232***<br>(0.012) | 1.261 |
| Female | -0.171***<br>(0.007) | 0.843 |
| Intercept | 1.777***<br>(0.005) |  |
| Observations | 2,739 |  |
| AIC | 7,713 |  |
| *Note:* | *p<0.1; **p<0.05; ***p<0.01 |  |

### Age-dependent survival and regional differences

The model including age, sex and region as covariates had the lowest DIC and was selected as the final model (Table 3). We extended the original model by estimating region-specific population growth rates (*λ_r_*) rather than a single shared *λ*, which improved the Bayesian p-value from 0.712 to 0.657, moving closer to the ideal value of 0.5. Parameter estimates for the final model are presented in Table 4 and estimated age-dependent survival probabilities are shown separately for males and females in Figure 2.

**Figure 2:**
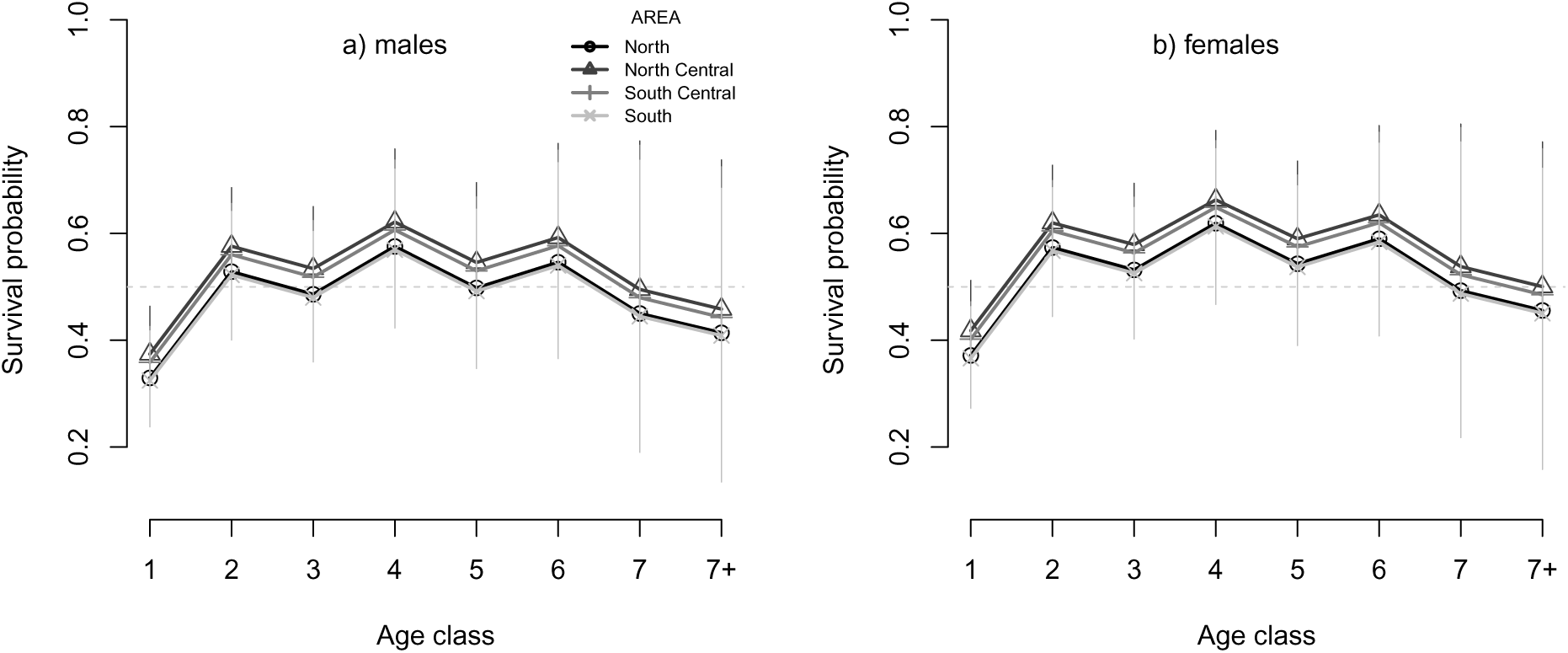
Estimated age-dependent survival of male and female red foxes in four different regions along a latitudinal gradient in Sweden. The dotted line indicates the 50% survival threshold. Survival was substantially lower for sub-adults (age class 1) than for adults across all regions and both sexes, producing a hump-shaped survival curve.

**Table 3:** Model selection and Bayesian p-values for estimates of age-dependent survival of red foxes in Sweden. Bayesian p-value for the final model reflects the improvement from region-specific growth rates.

| Model | Bayesian p-value | DIC |
| --- | --- | --- |
| Age + Sex + Region (region-specific $\lambda$) | 0.657 | 345.082 |
| Age + Sex + Region (shared $\lambda$) | 0.712 | 369.235 |
| Age + Sex | 0.544 | 372.144 |
| Age + Region | 0.025 | 400.950 |
| Age | 0.012 | 402.833 |
| Sex + Region | 0.000 | 496.588 |
| Sex | 0.000 | 501.042 |
| Region | 0.000 | 532.792 |
| Intercept | 0.000 | 536.054 |

**Table 4:** Posterior means and 95% credible intervals for the final age-dependent survival model with region-specific population growth rates. Parameters are on the logistic scale; age 1 and region N are reference categories fixed at zero. Age classes 7 and above are pooled into a single terminal class (7+). See appendix for full model specification.

| Parameter | Mean | Lower 2.5% | Upper 97.5% |
| --- | --- | --- | --- |
| Intercept | -0.717 | -1.061 | -0.354 |
| Age 2 | 0.832 | 0.545 | 1.149 |
| Age 3 | 0.660 | 0.387 | 0.968 |
| Age 4 | 1.029 | 0.631 | 1.522 |
| Age 5 | 0.709 | 0.296 | 1.193 |
| Age 6 | 0.909 | 0.351 | 1.627 |
| Age 7 | 0.503 | -0.595 | 1.706 |
| Age 7+ | 0.329 | -1.026 | 1.471 |
| Region NC | 0.196 | -0.348 | 0.731 |
| Region SC | 0.130 | -0.388 | 0.603 |
| Region S | -0.025 | -0.586 | 0.535 |
| Sex (female) | 0.186 | 0.116 | 0.263 |

Survival was substantially lower for sub-adults (age class 1) than for adults across all regions and both sexes. Survival increased to a peak around ages 3–6, then declined at older ages, producing a hump-shaped survival curve (Figure 2). The alternating pattern of estimates across adjacent age classes is a known mathematical property of the stable-stage back-calculation and does not reflect a biological signal (Skelly et al. 2023). Females had consistently higher survival than males across all regions and ages (logistic-scale coefficient: 0.186, 95% CrI: 0.116–0.263; Table 4).

Regional differences in survival were uncertain, with all region coefficients having 95% credible intervals spanning zero (Table 4). The pattern suggested slightly higher survival in the two central regions (NC and SC) relative to both the North and South, though this contrast was not credible. There was no simple latitudinal gradient in survival corresponding to the gradient observed in body weight.

### Regional population growth rates

Region-specific population growth rates are presented in Table 5. All four regions had wide credible intervals spanning *λ* = 1.0, indicating that the data do not provide strong evidence of systematic population growth or decline in any region during 1967–1971. However, the estimates suggest a latitudinal gradient, with the North showing the lowest mean growth rate (*λ* = 0.936, P(*λ >* 1) = 0.27) and South Central the highest (*λ* = 1.023, P(*λ >* 1) = 0.65). No pairwise contrast between regions was credible, as all 95% credible intervals for differences included zero.

**Table 5:**
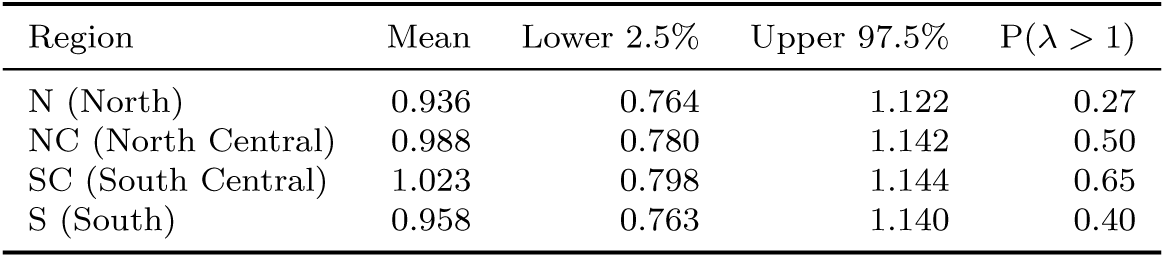
Posterior means and 95% credible intervals for region-specific population growth rates (*λr*) and the probability that *λ >* 1 (growing population) for red foxes in four latitudinal regions of Sweden, 1967–1971.

| Region | Mean | Lower 2.5% | Upper 97.5% | $P(\lambda > 1)$ |
| --- | --- | --- | --- | --- |
| N (North) | 0.936 | 0.764 | 1.122 | 0.27 |
| NC (North Central) | 0.988 | 0.780 | 1.142 | 0.50 |
| SC (South Central) | 1.023 | 0.798 | 1.144 | 0.65 |
| S (South) | 0.958 | 0.763 | 1.140 | 0.40 |

These patterns were broadly corroborated by independent regional harvest bag statistics for the same period. Harvest records showed a declining trend in the North (*λ* = 0.930), and relative stability or a slight increase in North Central (*λ* = 0.998), South Central (*λ* = 0.986), and South (*λ* = 1.025), consistent with the model’s lower *λ* estimate for the North relative to the other three regions.

### Age distribution

The relative age distributions in the different regions for each sex are presented in Figure 3. The distributions show the expected pattern of steeply declining proportions with increasing age class, with sub-adults (age 1) comprising approximately 54% of the harvest across all regions. The age distributions were broadly similar across regions and sexes, consistent with the modest and uncertain regional differences in survival estimated by the model.

**Figure 3:**
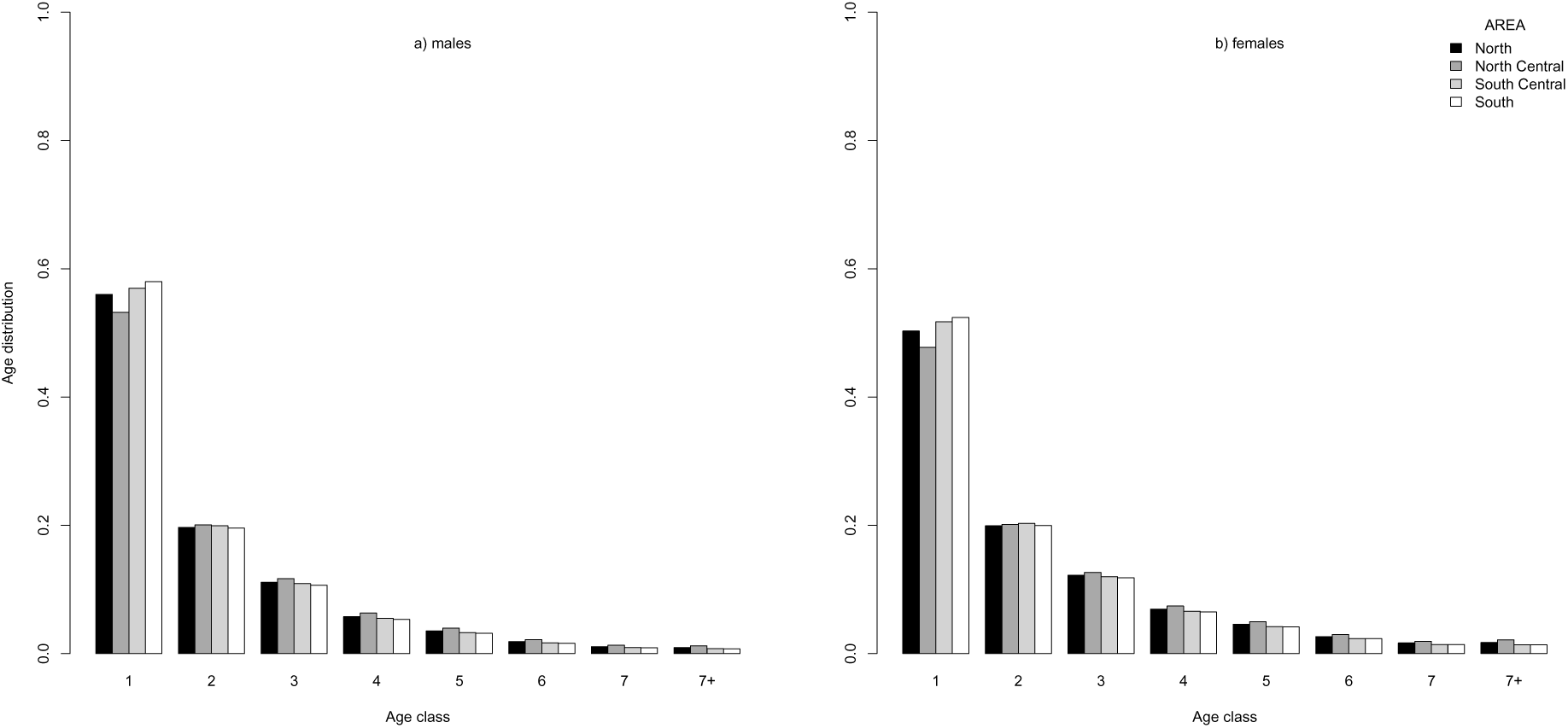
The estimated age distribution of male and female red foxes in four different regions in Sweden. The y-axis shows the proportion of foxes in each age class, and the x-axis shows the age classes (1 = sub-adults, 2–7+ = adults). The age distribution was broadly similar between regions and sexes, with a steep decline in the proportion of older age classes.

## Discussion

As expected, carcass weights of red foxes decreased with latitude, with southern foxes weighing approximately 1.27 times more than northern foxes. This pattern was consistent across both sexes and age classes, and the latitudinal effect on weight exceeded the sex difference. The decrease in weight with latitude is likely explained by a reduced availability of garbage, food waste, offal remains and prey availability, in combination with harsher winter conditions in the northern regions, which can limit growth and body size. Because body mass and body size are closely correlated in mammals, we interpret the latitudinal weight differences reported here as indicative of broader differences in body size, and discuss them in that context below. The trend in body size follows the regional productivity gradient, as has been shown for several medium-sized carnivores (Yom-Tov, Yom-Tov, and Zachos, 2013; Wereszczuk et al. 2021). Diet and human subsidies are more important than climate for the development of body size in productive habitats in Spain and Denmark (Yom-Tov, Yom-Tov, Barreiro, et al. 2007; Gortázar et al. 2000). Nutritional conditions during early development affect final body size, which can in turn affect both optimization of life histories and population dynamics (Lindström, 1999; Bartoń and Zalewski, 2007; Reeves et al. 2020). Body size of red fox females probably affects the size of the offspring, similar to that of red deer (*Cervus elaphus*) females, which, if born in poor years, could not respond to good adult conditions the way well-grown females can (Nussey, Clutton-Brock, Albon, et al. 2005; Nussey, Clutton-Brock, Elston, et al. 2005). In addition, female territoriality and high density in the southernmost region probably force subordinate females to abstain from breeding (Macdonald, 1979; von Schantz, 1984). The ones that do breed are probably the ones in better condition, and the size of their offspring can be expected to be larger than the offspring of the subordinate females if they had produced young. However, territoriality is not a species specific characteristic, and it is likely not an economically defensible behavior in the northern regions with more scarce and unpredictable food resources (Powell, 2000). Therefore, we expect an increasing degree of scramble and decreasing degree of contest competition with increasing latitude. Thus, the weight distribution of the harvested foxes in the southernmost region is probably skewed towards heavier individuals, and reinforces the difference to other regions. This pattern is contrary to Bergmann’s rule, which predicts larger body sizes at higher latitudes, but is consistent with other studies that have found exceptions to this rule in carnivores (Frafjord and Stevy, 1998; Meiri, Guy, et al. 2009; Yom-Tov, Yom-Tov, Barreiro, et al. 2007).

Males were approximately 20% heavier than females across all regions and age classes. This sexual dimorphism in body size is common in many carnivores and can be attributed to the differences in reproductive strategies (Gittleman, 1985; Moors, 1980). We also found a small but significant increase in weight with age among adults, with foxes gaining approximately 1.1% in weight per year. This suggests that individuals that survive to older ages may continue to grow larger, which could have implications for survival and reproductive success. Therefore, it is important to consider age structure when interpreting patterns of body size variation, as changes in the age distribution could affect the average body size observed in different regions as suggested by Theriot et al. (2024). In a study of intensive hunting of red foxes, it was shown that the mean longevity decreased, and age structure of the population changed to a higher proportion of younger individuals in the population (Yoneda and Maekawa, 1982). However, in our study, the age distributions were broadly similar across regions, suggesting that the observed latitudinal gradient in weight was not primarily driven by differences in age structure.

The regional lambda estimates from the age-at-harvest model were independently corroborated by the harvest bag statistics. The northern decline and southern growth gradient visible in the harvest data was consistent with the latitudinal productivity gradient, with northern regions showing lower fox population growth rates during this period. Males showed consistently lower survival than females across all regions and ages, and lower survival of males compared to females has been reported by Heydon and Reynolds (2000), although at lower survival rates. A male-biased sex ratio in the harvest is common in samples of red foxes (Englund, 1970; Yoneda and Maekawa, 1982). Harvest is an important cause of mortality in red foxes, and males may be more vulnerable to harvest due to larger home ranges and more exploratory behavior (Heydon and Reynolds, 2000). The estimates of age-dependent survival are similar to earlier studies of red foxes in Sweden (Devenish-Nelson et al. 2013; Willebrand et al. 2022), with low survival for sub-adults and a hump-shaped curve for adults. The highest survival was recorded in the central regions, but the estimates of age-dependent survival did not show a clear latitudinal gradient. The regional differences were uncertain, with all credible intervals spanning zero.

The lack of a clear latitudinal pattern in survival corresponding to the strong gradient observed in body weight suggests that density-dependent mortality may play an equalizing role across regions. Higher productivity in the south likely supports both heavier foxes and higher population densities, with the resulting intraspecific competition potentially suppressing survival to levels similar to the less productive northern regions. If recruitment of young also varies among regions in proportion to productivity, the overall population growth rates may be more similar across the latitudinal gradient than the large differences in body condition would suggest, because the *realized* recruitment into the breeding population end up similar across regions. This is analogous to work on large herbivores showing that recruitment can vary substantially across populations while adult survival remains relatively constant, resulting in similar population growth rates despite differences in productivity and body condition (Gaillard, Festa-Bianchet, et al. 2000).

### Assumptions, limitations and future directions

The age-at-harvest model used to estimate survival in this study rests on two key assumptions that deserve consideration. First, the model assumes that all age classes are equally vulnerable to harvest, but harvest is an important source of mortality in red foxes and selectivity may vary with age, sex, and region. Males, with larger home ranges and more exploratory behaviour, may be more vulnerable to harvest than females, which could bias the estimated sex difference in survival. Similarly, if sub-adults are more vulnerable to harvest than adults, the low sub-adult survival estimates may partly reflect harvest selectivity rather than true survival differences. A concurrent analysis of 126 radio-collared red foxes from the same study system has provided independent survival estimates that corroborate the robustness of our age-at-harvest results and that harvest is the dominating mortality cause (Willebrand et al., unpublished).

Second, the model assumes a stable age structure, meaning that the relative proportions of each age class in the harvest reflect the true age structure of the population. This assumption is most likely violated during periods of rapid population change, such as following disease outbreaks or major environmental perturbations. Our data were collected between 1967 and 1971, prior to the first outbreak of sarcoptic mange in Sweden in the late 1970s, and harvest records suggest that populations were broadly stable during this period, particularly in the southern and central regions. However, the northern region showed signs of decline during the study period, as reflected in both the harvest bag statistics and the estimated population growth rate (*λ* = 0.936), which suggests that the stable age structure assumption may be least well satisfied in the North. The use of region-specific population growth rates partially relaxes this assumption by allowing the age structure to deviate from stability in proportion to *λ_r_*, which is one reason we prefer the region-specific model over the shared-*λ* formulation.

Dispersal represents a further potential source of bias in the age-at-harvest model, as immigration of young foxes from the more productive southern regions into the north could inflate the proportion of young individuals in northern harvest samples, potentially biasing survival estimates upward in those regions. Long-distance dispersal has been documented in red foxes in Sweden, with individuals moving several hundred kilometers between natal and settlement areas (Walton et al. 2018), making inter-regional movement biologically plausible. However, if dispersing individuals are predominantly young subadults, as is typical in carnivores, this would primarily affect sub-adult survival estimates rather than adult survival, and the direction of the bias would be consistent across regions rather than generating the uncertain regional pattern we observed.

## Conclusions

Our results demonstrate a clear latitudinal gradient in body size of red foxes in Sweden, with southern foxes weighing approximately 1.27 times more than northern foxes. This pattern was consistent across both sexes and age classes, with the latitudinal effect exceeding the sex difference in body size, and is contrary to Bergmann’s rule. The decrease in body size with latitude is consistent with a productivity-driven resource limitation, where reduced prey availability and harsher winter conditions in the north limit growth and body size during development. Despite the strong latitudinal gradient in body size, survival did not show a corresponding pattern — regional differences in survival were uncertain, with all credible intervals spanning zero. The decoupling of body condition and survival across regions suggests that factors beyond individual nutritional state regulate mortality similarly across the latitudinal gradient. Density-dependent processes are a plausible mechanism, whereby higher productivity in the south supports both heavier foxes and higher population densities, potentially suppressing survival through intraspecific competition to levels similar to the less productive northern regions. This hypothesis remains to be tested with concurrent density estimates across the latitudinal gradient. Regional population growth rates, broadly corroborated by harvest bag statistics, were consistent with a declining trend in the north and relative stability across the central and southern regions, reflecting the underlying productivity gradient and suggesting that body condition and population dynamics are coupled at the regional scale despite the absence of a clear survival gradient.

## Acknowledgements

We are grateful to the dedication and persistent efforts of the late Professor Jan Englund. The Swedish Environmental Protection Board, the Swedish Hunters Association, and the University of Inland Norway provided funding for the study.

## Author contributions

Tomas Willebrand: Conceptualization (equal); Data curation (lead); Formal analysis (lead); Funding acquisition (equal); Investigation (equal); Methodology (lead); Project administration (lead); Writing – original draft (lead); Writing – review and editing (lead). Gustaf Samelius: Conceptualization (equal); Formal analysis (equal); Investigation (equal); Methodology (equal); Validation (equal); Writing – review and editing (equal). Zea Walton: Conceptualization (equal); Formal analysis (equal); Investigation (equal); Methodology (equal); Validation (equal); Writing – review and editing (equal). Morten Odden: Conceptualization (equal); Investigation (equal); Methodology (equal); Validation (equal); Writing – review and editing (equal). Jan Englund: Provided data; Conceptualization (equal).

## Data availability

Data on body weight and age distribution of harvested foxes in different regions are available at Zenodo.org (doi:10.5281/zenodo.21023047).

## Appendix

## Jags models

*Model use to estimate age-dependent survival:*

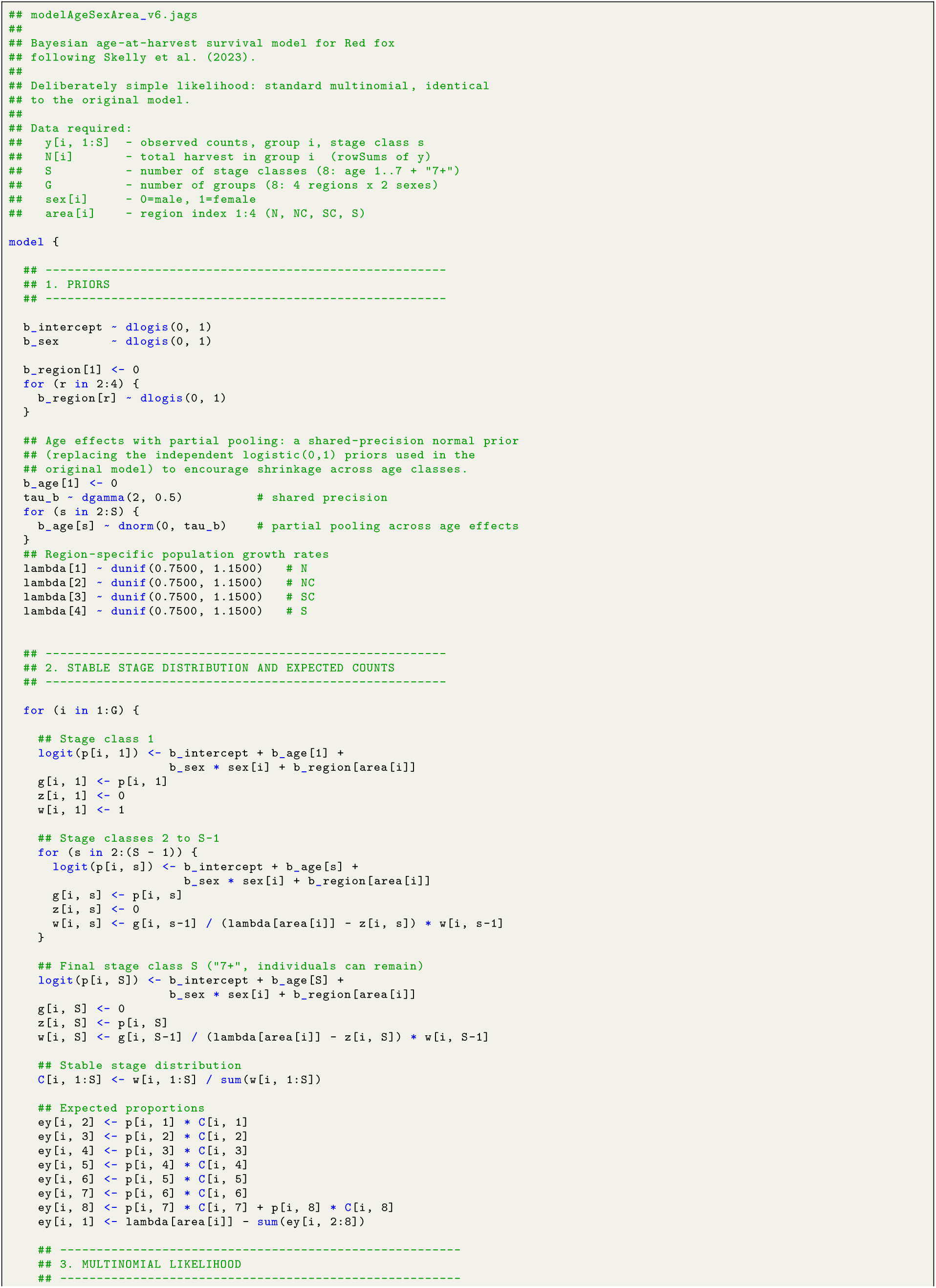
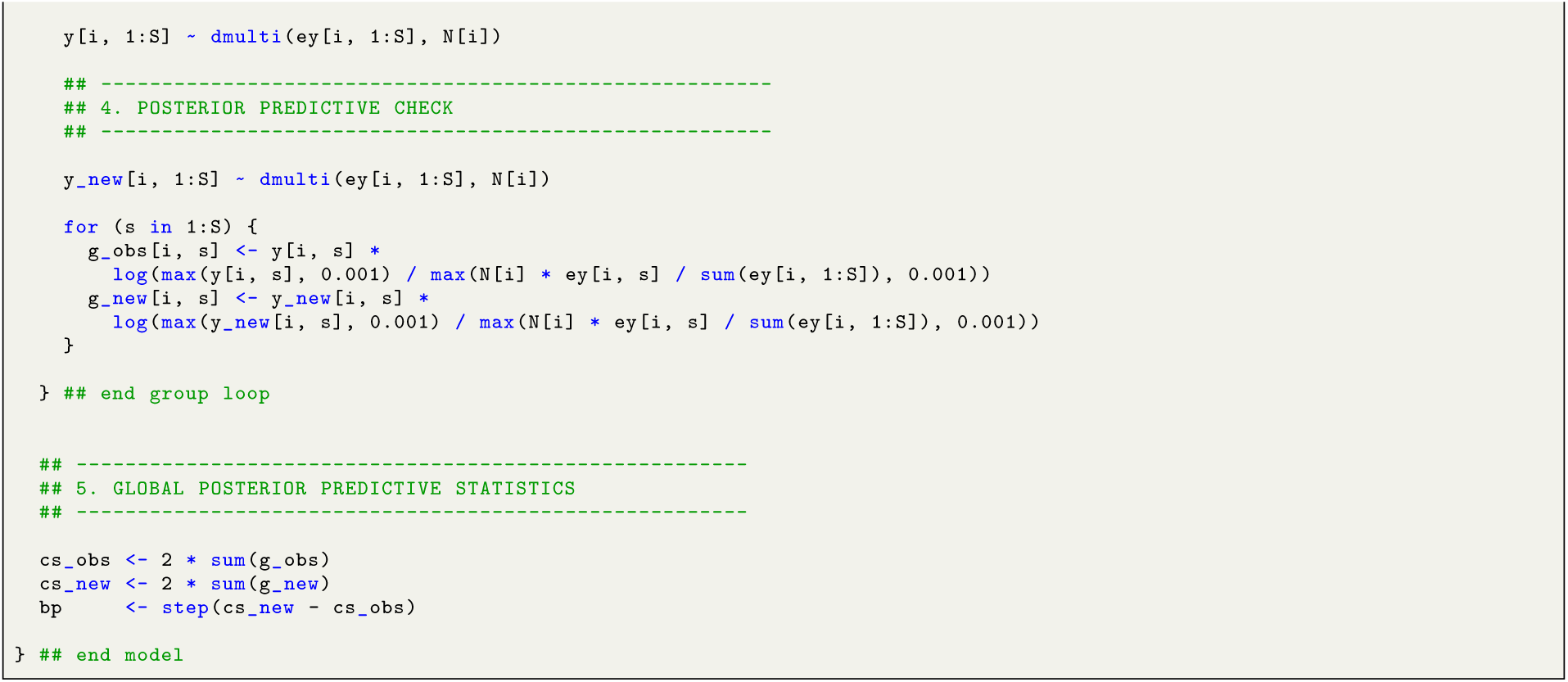

